# SARS-CoV-2 genomic and quasispecies analyses in cancer patients reveal relaxed intrahost virus evolution

**DOI:** 10.1101/2020.08.26.267831

**Authors:** Juliana D. Siqueira, Livia R. Goes, Brunna M. Alves, Pedro S. de Carvalho, Claudia Cicala, James Arthos, João P.B. Viola, Andréia C. de Melo, Marcelo A. Soares, on behalf of the INCA COVID-19 Task Force

**Author notes:** Corresponding author (JDS). Authors contributed equally to this work (Juliana D. Siqueira, Livia R. Goes, Brunna M. Alves). Membership of INCA COVID-19 Task Force is provided in the Acknowledgments.

## Abstract

Numerous factors have been identified to influence susceptibility to SARS-CoV-2 infection and disease severity. Cancer patients are more prone to clinically evolve to more severe COVID-19 conditions, but the determinants of such a more severe outcome remain largely unknown. We have determined the full-length SARS-CoV-2 genomic sequences of cancer patients and healthcare workers (HCW; non-cancer controls) by deep sequencing and investigated the within-host viral quasispecies of each infection, quantifying intrahost genetic diversity. Naso- and oropharyngeal SARS-CoV-2^+^ swabs from 57 cancer patients and 14 healthcare workers (HCW) from the Brazilian Cancer Institute were collected in April–May 2020. Complete genome amplification using ARTIC network V3 multiplex primers was performed followed by next-generation sequencing. Assemblies were conducted in Geneious R11, where consensus sequences were extracted and intrahost single nucleotide variants (iSNVs) were identified. Maximum likelihood phylogenetic analysis was performed using PhyMLv.3.0 and lineages were classified using Pangolin and CoV-GLUE. Phylogenetic analysis showed that all but one strain belonged to clade B1.1. Four genetically linked mutations known as the globally dominant SARS-CoV-2 haplotype (C241T, C3037T, C14408T and A23403G) were found in the majority of consensus sequences. SNV signatures of previously characterized Brazilian genomes were also observed in most samples. Another 85 SNVs were found at a lower frequency (1.4-19.7%). Cancer patients displayed a significantly higher intrahost viral genetic diversity compared to HCW (p = 0.009). Intrahost genetic diversity in cancer patients was independent of SARS-CoV-2 Ct values, and was not associated with disease severity, use of corticosteroids, or use of antivirals, characteristics that could influence viral diversity. Such a feature may explain, at least in part, the more adverse outcomes to which cancer/COVID-19 patients experience.

**Author Summary:** Cancer patients are more prone to clinically evolve to more severe COVID-19 conditions, but the determinants of such a more severe outcome remain largely unknown. In this study, phylogenetic and variation analysis of SARS-CoV-2 genomes from cancer patients and non-cancer healthcare workers at the Brazilian National Cancer Institute were characterized by deep sequencing. Viral genomes showed signatures characteristic of Brazilian viruses, consistent with the hypothesis of local, community transmission rather than virus importation from abroad. Despite most genomes in patients and healthcare workers belonging to the same lineage, intrahost variability was higher in cancer patients when compared to non-cancer counterparts. The intrahost genomic diversity analysis presented in our study highlights the relaxed evolution of SARS-CoV-2 in a vulnerable population of cancer patients. The high number of minor variations can result in the selection of immune escape variants, resistance to potential drugs, and/or increased pathogenicity. The impact of this higher intrahost variability over time warrants further investigation.

## Introduction

In December 2019, a new form of pneumonia was described in patients with severe acute respiratory syndrome in the city of Wuhan, province of Hubei, China [1]. Soon after, a new beta-coronavirus was identified as the causative agent of that disease [2]. The new virus was named Severe Acute Respiratory Syndrome Coronavirus 2 (SARS-CoV-2), and the disease was called Coronavirus Disease 2019 (COVID-19) [2]. Since its initial discovery, COVID-19 has become a pandemic of catastrophic proportions, with over 17 million confirmed cases of viral infection and over 670,000 deaths worldwide (https://www.worldometers.info/coronavirus/, last accessed on July 30^th^, 2020).

Numerous demographic, clinical, genetic, and behavioral factors have been identified to influence susceptibility to SARS-CoV-2 infection and, among those infected, the severity of the disease, including the risk of death. Those factors include age, sex [3], genetic loci of certain cytokines/chemokines and the ABO blood system group [4, 5], smoking history [6], obesity and underlying comorbidities such diabetes, hypertension, lung diseases [7, 8], and cancer [9–11]. Among cancer patients, those with malignancies of hematological origin have been reported as particularly vulnerable to COVID-19 [12].

SARS-CoV-2 is a single-stranded RNA virus that replicates using an RNA-dependent RNA polymerase. As such, the virus is subjected to high rates of nucleotide sequence changes, and has evolved through molecular evolution and founder effects during its explosive spread throughout the globe. Virus replication rates directly impact the accumulation of mutations in the virus genome, enabling the existence of a viral quasispecies (a swarm of different, yet highly related, viral entities) within an infected host. Although within-host variations of SARS-CoV-2 have been documented [13, 14], the impact of underlying comorbidities that promote persistent viral RNA detection and shedding on virus evolution remains to be elucidated. Moreover, viral genetic variation, as a source of novel mutations, may hinder future therapeutic antiviral and vaccine strategies targeting COVID-19, by the selection of drug-resistant and vaccine escape mutants [15].

In the present work, we have determined the full-length SARS-CoV-2 genomic sequences of 57 cancer patients and 14 healthcare workers (non-cancer controls) employing next-generation sequencing (NGS), and analyzed their epidemiological relatedness and lineage classification. This approach also allowed us to study the within-host viral quasispecies of each infection, quantify intrahost viral genetic diversity and characterize specific genetic changes with potential to impact the virus biology. Finally, we have also assessed associations between viral diversity and patients’ clinical and laboratory characteristics, thereby identifying determinant factors of viral evolution in this particular group of patients.

## Results

### Clinical characteristics of the studied population

Summarized demographic and clinical characteristics of the patients and healthcare workers from whom SARS-CoV-2 sequences were studied can be seen in Table 1. Among patients, the median age was 61 years and most of them (72%) had solid malignancies, 16% of patients used corticosteroids and 14% used oseltamivir previously or during COVID-19 diagnosis specimen collection. Among healthcare workers (HCW), the median age was 40 years and most (86%) were female. The most prevalent COVID-19 symptoms among patients were cough, fever and dyspnea. Death from COVID-19 occurred in 33.3% of the cases. For HCW, cough and coryza were the mainly reported COVID-19 symptoms (85.7% each), and all subjects recovered from the disease, with no deaths reported. No difference was found in sex distribution between the two groups (p = 0.118), but HCW had a lower median age when compared to cancer patients (p < 0.001).

**Table 1.** Demographic and clinic characteristics of the cancer patients and healthcare workers studied.

| Characteristic | Patients (%)<br>n = 57 | Healthcare workers (%)<br>n = 14 |
| --- | --- | --- |
| Age years (median, range) | 61 (9-79) | 40.5 (33-57) |
| Age |  |  |
| < 25 years | 3.5 | 0 |
| 25 to 64 years | 57.9 | 100 |
| ≥ 65 years | 38.6 | 0 |
| Gender |  |  |
| Female | 61.4 | 85.7 |
| Male | 38.6 | 14.3 |
| Symptoms at COVID-19 diagnosis |  |  |
| Cough | 59.6 | 85.7 |
| Fever | 57.9 | 57.1 |
| Dyspnea | 56.1 | 7.1 |
| Fatigue | 24.6 | 21.4 |
| Diarrhea | 14.0 | 7.1 |
| Nausea/Vomiting | 12.3 | 0 |
| Anorexia | 7.0 | 0 |
| Sore throat | 5.3 | 42.8 |
| Myalgia | 3.5 | 0 |
| Headache | 3.5 | 42.8 |
| Anosmia | 3.5 | 42.8 |
| Ageusia | 3.5 | 0 |
| Coryza | 3.5 | 85.7 |
| None | 0 | 0 |
| Missing | 7.0 | 0 |
| Death |  |  |
| Yes, from COVID-19 | 33.3 | 0 |
| Yes, other cause | 5.3 | 0 |
| No | 54.4 | 100 |
| Missing | 7.0 | 0 |
| Smoking |  |  |
| Past/current | 21.0 | NC* |
| Never | 24.6 | NC |
| Missing | 54.4 | NC |
| Primary cancer site |  |  |
| Solid Tumors | 71.9 | NA<sup>#</sup> |
| Hematological malignancies | 28.1 | NA |
| Metastatic disease |  |  |
| Yes, to the lung | 14.0 | NA |
| Yes, to other organs | 24.6 | NA |
| No | 47.4 | NA |
| Missing | 14.0 | NA |
| Use of corticosteroid |  |  |
| Yes | 19.3 | NA |
| No | 87.2 | NA |
| Missing | 3.5 | NA |
| Use of oseltamivir |  |  |
| Yes | 14.0 | NA |
| No | 82.4 | NA |
| Missing | 3.5 | NA |

\*NC – not collected; # NA – not applicable

### Sequence coverage, quality and metrics

A total of 27,433,528 reads were obtained from sequencing, with an average of 382,118 reads per sample, ranging from 217,922 to 631,796 reads. Reads of each sample were assembled with Wuhan-Hu-1 reference genome with a minimum mapping quality of 30 Phred and the average depth coverage obtained was 1,468 (465-2,530). The coverage was heterogeneous across the genome but was similar among the samples (S1 Fig). Consensus sequences containing more than 97.9% of the SARS-CoV-2 complete genome were generated from all 57 cancer patient and 14 HCW samples.

### Phylogenetic and epidemiological profile of SARS-CoV-2 sequences

SARS-CoV-2 genome sequence submission to the *Pangolin* and *CoV-GLUE* algorithms resulted in the same lineage classification in all cases, defining all but one virus belonging to clade B1.1, while the remaining sequence was classified as B.1. A phylogenetic analysis of the viruses together with sequences previously defined as Brazilian circulating strains B1.1-BR and B1.1-EU/BR showed that most B1.1 genomes generated in this study clustered with B1.1-BR sequences (Fig 1A) [16]. A phylogenetic tree including all local SARS-CoV-2 sequences isolated from patients residing in the state of Rio de Janeiro available at the GISAID database (accessed on July 27th, 2020, S1 Table) was performed to investigate potential epidemiological linkage between samples (Fig 1B). We noted that some of the viruses sequenced at INCA clustered in clades containing identical sequences, suggesting a transmission link between the study subjects. In some instances, both cancer patients and HCW were involved in those epidemiological clusters. Although in some cases sequences from outside the hospital were also identical to viruses from our series, therefore not excluding the possibility of community transmission, the most likely scenario for those cases is a nosocomial transmission between patients and/or HCW.

**Fig 1.**
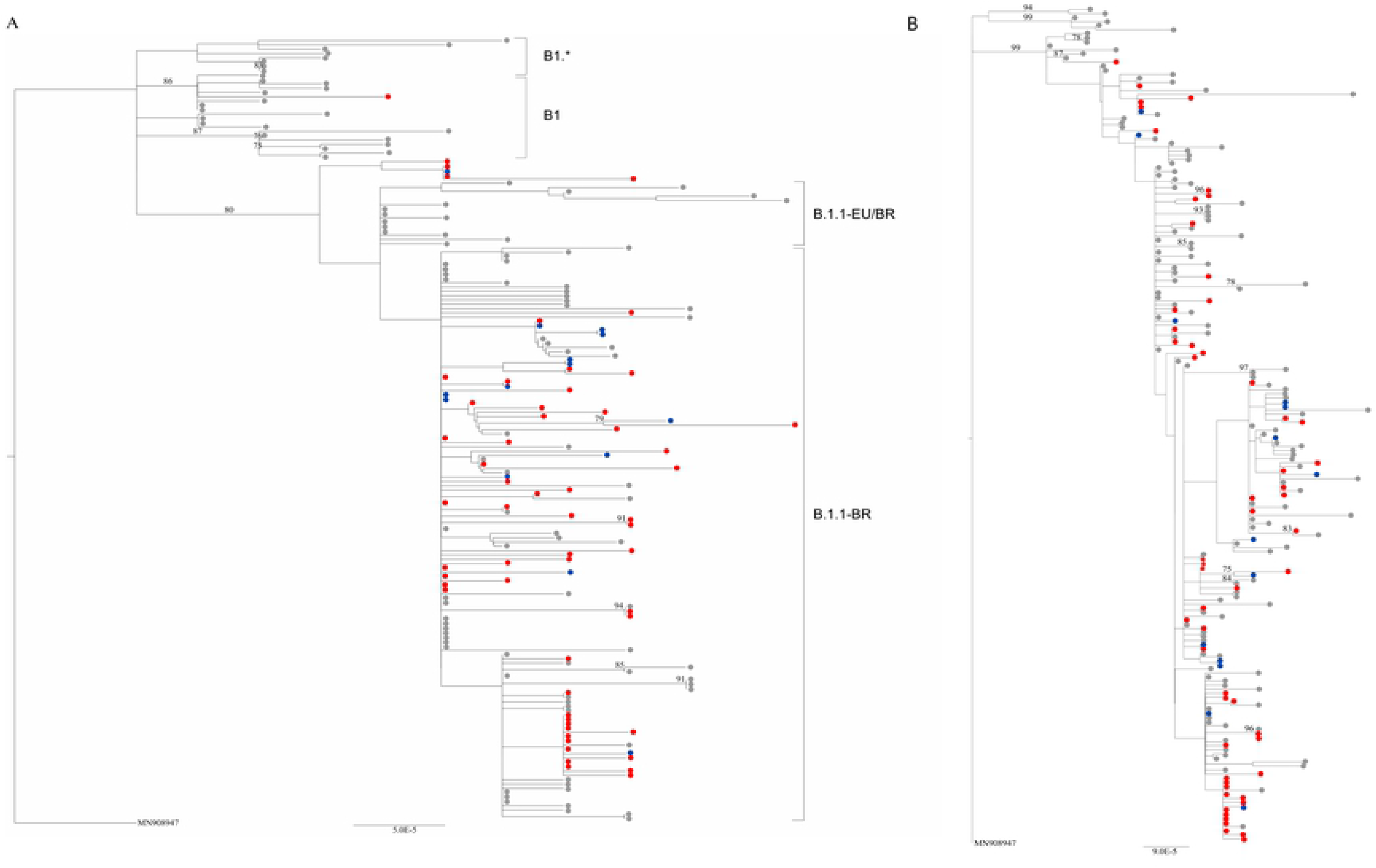
Maximum likelihood phylogenetic trees of near full-length SARS-CoV-2 genomes characterized. Tree including cancer patients (depicted in red circles), healthcare workers (in blue), Brazilian sequences classified as B1, B1.* and the Brazilian circulating strains B1.1-BR/ B.1.1-EU/BR available on GISAID (in gray). (B) Tree showing epidemiological linkage of cancer patients (shown in red), healthcare workers (in blue) and all SARS-Cov-2 sequences from Rio de Janeiro state (in gray) available on GISAID Database. In both cases, GISAID was accessed on July 27th, 2020. Bootstrap values greater than 70 are shown in both trees.

### Single nucleotide variations across the SARS-CoV-2 genomes

Overall, 95 single nucleotide variations (SNVs) and three deletions were found across the SARS-CoV-2 genomes analyzed (Fig 2). Four genetically linked mutations previously described as the globally dominant haplotype in April 2020 were found in the majority of our consensus sequences: C241T (100%; 5’UTR region), C3037T (98.6%; silent mutation), C14408T (100%; resulting in P4715L/P323L amino acid change in ORF1ab) and A23403G (100%; resulting in D614G amino acid change in S) [17]. Additionally, SNV signatures of previously characterized Brazilian genomes were found in most samples, such as G28881A and G28882A (98.6%; resulting in R203K change in N), G28883C (98.6%; resulting in G204R change in N), T27299C (91.6%; resulting in I33T change in ORF6), and T29148C (90.1%; resulting in I292T change in N) [16, 18]. The two latter SNVs are synapomorphic traits of the B1.1-EU/BR and B1.1-BR Brazilian circulating strains [16]. Another 85 SNVs were observed in our sequences in a lower frequency (1.4-19.7%; S2 Table), including nine non-synonymous mutations in S protein (V16F, V367L, K558N, Q675H, A879V, S939F, V1176F, K1191N and G1219V). Deletions were found in three genomes: a 12-bp in-frame deletion in S (comprising positions 21603-21614), a 6-pb in-frame deletion in ORF3a (25710-25715) and a 244-pb frameshift deletion in ORF7 (27508-27751), resulting in a truncated protein. All deletions were confirmed by Sanger sequencing (data not shown).

**Fig 2.**
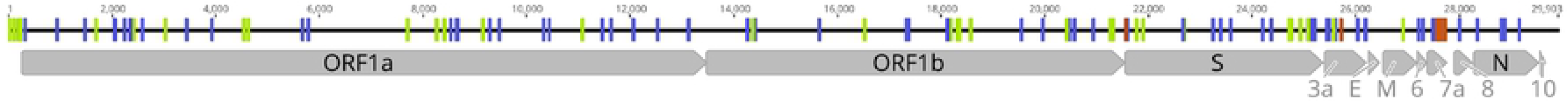
Distribution of variations in the SARS-CoV-2 genome. Variations found across different ORFs of SARS-CoV-2 genomes analyzed. Synonymous mutations and changes found in noncoding regions are highlighted in green, non-synonymous mutations in blue and deletions in red. Genome coordinates are relative to the SARS-CoV-2 Wuhan-Hu-1 reference sequence genome (GenBank acc.# MN908947).

### SARS-CoV-2 intrahost genetic diversity

The next-generation sequencing method used for the study viruses allowed us to assess the intrahost SNVs (iSNVs) that compose each subject’s viral quasispecies. The number of iSNVs across the viral genome can be visualized in Figs 3A (patients) and 3B (HCW). All but one iSNV with intrahost frequency greater than 20% were found exclusively in cancer patients’ samples (Fig 4). Interestingly, patients displayed a significantly higher intrahost viral genetic diversity when compared to HCW (p = 0.009; Fig 5A) and remained significant even after outlier subjects with higher virus diversity were excluded from the analysis (p = 0.029; Fig 5B). Viral genetic diversity within each ORF was compared between the two groups, and cancer patients carried a higher genetic diversity in ORF 1A when compared to non-cancer HCW (p = 0.045).

**Fig 3.**
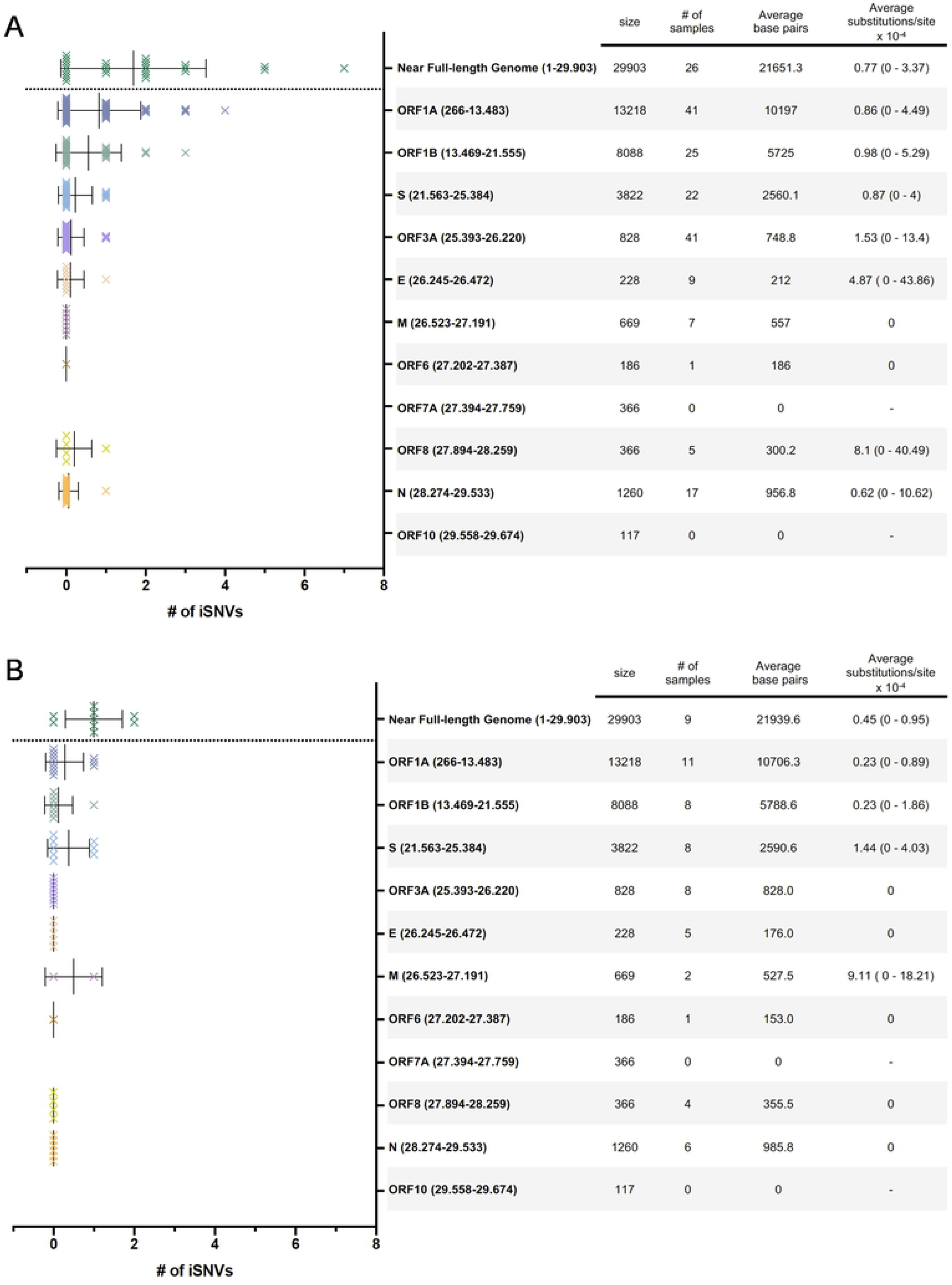
Number of intrahost single nucleotide variants (iSNVs) per ORF analyzed. Data for cancer patients (A) and healthcare workers (B) are shown. iSNV shown are those with an intrahost frequency greater than 2% and a minimum depth coverage of 500x. The table on the right shows ORF names and genome coordinates based on the SARS-CoV-2 Wuhan-Hu-1 reference sequence genome (GenBank acc.# MN908947), ORF size in bp, number of samples analyzed that fulfilled the criteria, average base pairs analyzed (considering a minimum depth coverage of 500x for at least 60% of the ORF region) and average (min - max) substitutions per site × 10^−4^.

**Fig 4.**
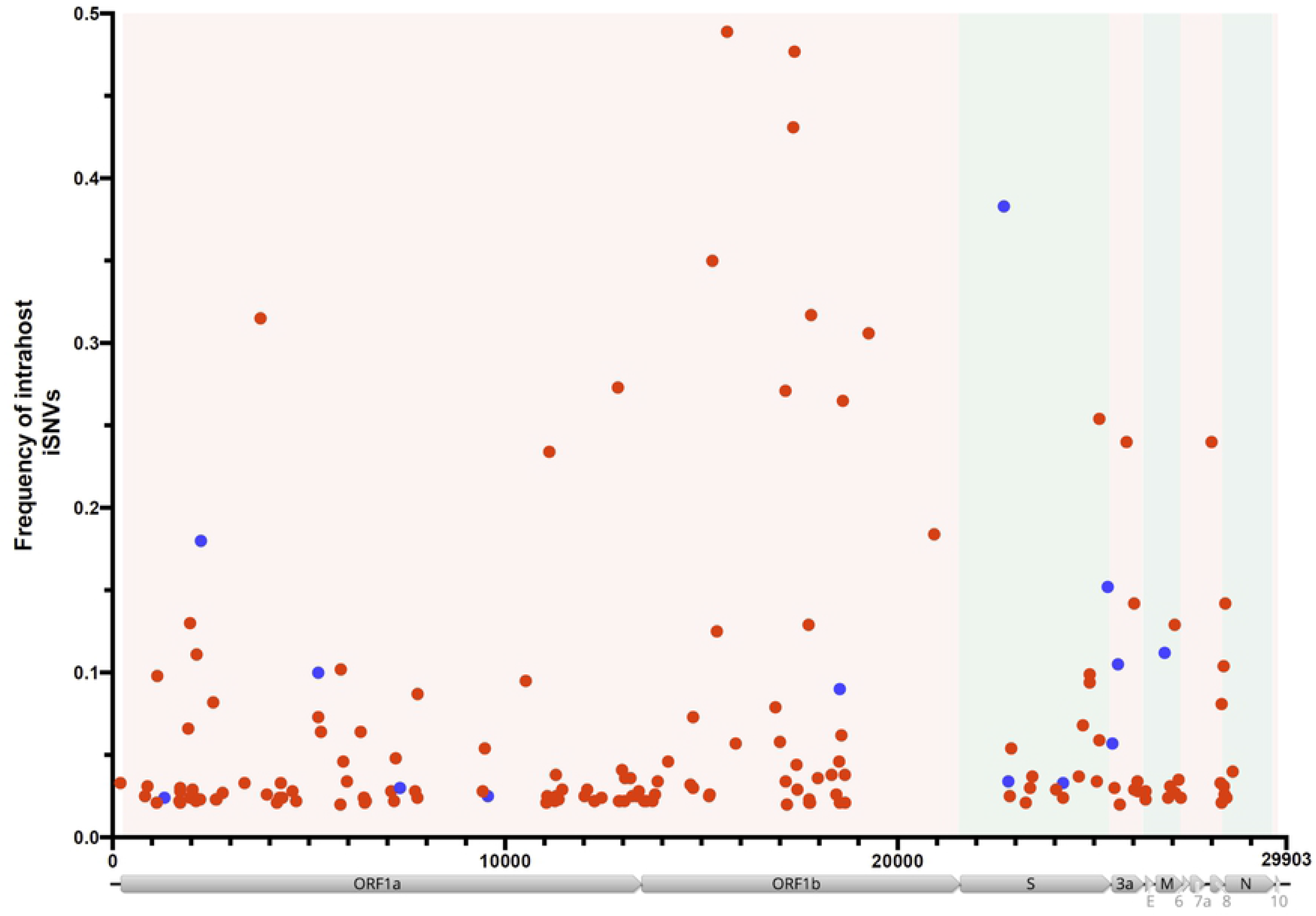
Frequency of intrahost single nucleotide variants (iSNVs) across the SARS-CoV-2 genome. Distribution (x-axis) and frequency (y-axis) of iSNVs (>2%) with a minimum depth coverage of 500x. Red dots represent cancer patients and blue dots represent healthcare worker samples. Structural genes (S, E, M and N) are highlighted in green and non-structural genes in light-red. Genome coordinates are relative to the SARS-CoV-2 Wuhan-Hu-1 reference sequence genome (GenBank acc.# MN908947).

**Fig 5.**
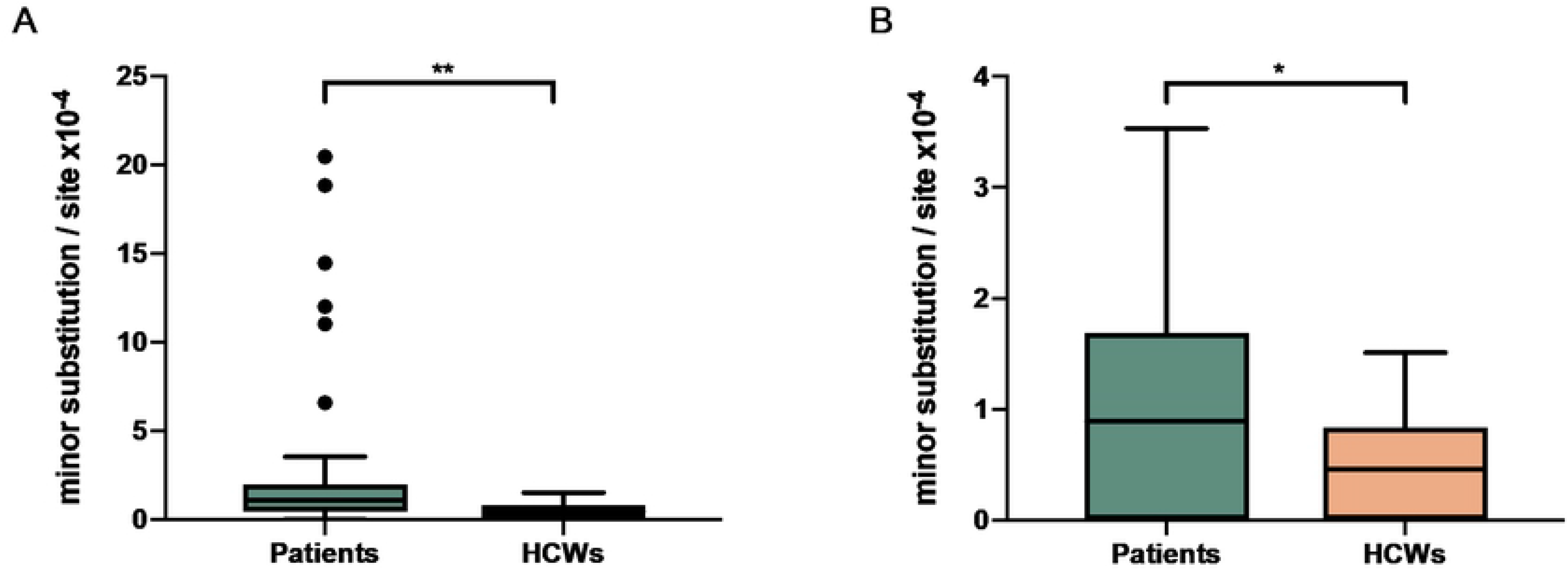
Viral genetic diversity in cancer patients and healthcare workers (HCWs). Diversity was calculated by number of minor substitutions per site x10^−4^. Tukey boxplots show the viral genetic diversity in cancer patients (n=57) compared to HCW (n=14) (Mann-Whitney test (two-tailed), **p=0.0093) (A). The difference remains significant when outlier patients (n=6) are removed from the analysis (Mann-Whitney test (two-tailed), *p=0.0299) (B).

As the within-host genetic diversity of viruses is commonly associated with viral replication, we have evaluated the correlation of the quasispecies diversity in our subjects with the Ct values obtained in the RT-PCR swab tests of the same samples. Ct values work as a proxy for SARS-CoV-2 viral load in samples and are expected to be inversely correlated with viral diversity and replication. Surprisingly, however, Ct of the samples did not inversely correlate with viral diversity, but rather showed a positive significant correlation, despite having r_s_ values below 0.5 (Fig 5). This was true for patients’ samples (r_s_ = 0.490; p = 0.001; Fig 6A) and also when all patients and HCW were analyzed together (r_s_ = 0.478; p < 0.0001; Fig 6B). Of note, no differences were found when Ct values were compared between the two groups (p = 0.175). Despite the above mentioned age difference observed between HCW and cancer patients, age did not correlate with viral genetic diversity (S2 Fig, p = 0.844).

**Fig 6.**
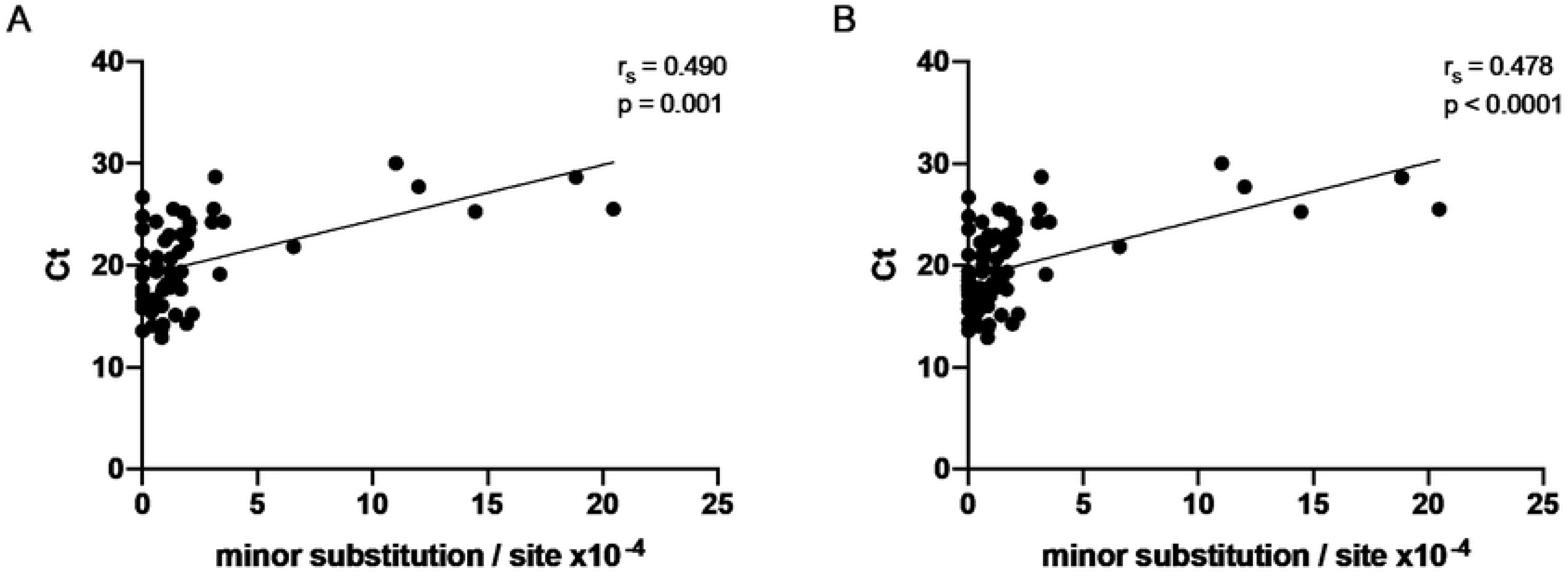
Spearman correlation of viral genetic diversity and Ct values. Cancer patients samples (n=57) showed a positive correlation with Ct values (A). The same result was found when both patients and healthcare workers samples were combined (n=71) (B). Spearman correlation analysis r_s_ and p-values are indicated.

Regarding patients’ characteristics, intrahost virus diversity was not associated with disease severity (overall death [p = 0.632] or death due to COVID-19 [p = 0.934], ICU requirement [p = 0.612]), use of corticosteroids chronically or during COVID-19 course (p = 0.333), or use of oseltamivir prior to COVID-19 diagnosis (p = 0.144; S3A-E Fig). We also assessed the potential association of cancer patients with hematological malignancies compared to those with solid cancers, but no association was found (p = 0.473; S3F Fig).

## Discussion

The biology of SARS-CoV-2 infection in humans is striking to infectious disease clinicians worldwide, because no viral infection has been previously seen with such an enormous range of phenotypic outcomes, from no symptoms to severe respiratory distress and death. Most of this physiological variance, however, has been attributed to host genetic and behavioral factors. Numerous characteristics have been associated with susceptibility to SARS-CoV-2 infection and disease severity among infected subjects, and underlying comorbidities seem to play a major role in unfavorable disease outcomes. Chronic non-communicable diseases such as cancer are among those conditions. Cancer patients have been reported to be more prone to SARS-CoV-2 infection and to clinically evolve to more severe conditions upon infection [9–11], but the determinants of these severe outcomes remain largely unknown.

In this study we have evaluated the near full-length sequences of SARS-CoV-2-infected cancer inpatients in one of the largest public cancer hospitals in South America, the Brazilian National Cancer Institute, and compared these sequences with those generated from healthcare professionals from the same institution. These complete SARS-CoV-2 genomes showed signatures characteristic of the virus that spread globally and is currently the predominant strain [17]. All but one virus also belonged to clade B1.1, which is the clade primarily circulating in the Americas. The viral genomes also displayed sequence features of other already characterized Brazilian viruses, consistent with the hypothesis of local, community transmission rather than virus importation from abroad. In fact, the timeframe of the analyzed infections (from April 7^th^ to May 5^th^, 2020) is consistent with a period in Brazil where community virus transmission was already established and ongoing [18]. Moreover, as a public and free hospital in the Brazilian Public Health System, INCA is also likely to admit patients with low socioeconomic resources who are mostly unable to travel abroad and most likely acquired viral infections from local sources.

We explored the evolutionary and phylogenetic relationships between the SARS-CoV-2 sequences of the studied samples. Upon a phylogenetic inference with viral sequences isolated from other infected subjects residing in the state of Rio de Janeiro (the same geographic location of the study site), we found that almost half of the sequences from our subjects lie in clusters with sequences from other patients and/or from HCW. Some of the consensus sequences within each cluster were identical, suggesting a direct epidemiological link between those groups of patients/HCW. Some identical sequences retrieved from the database representing subjects from the community outside the hospital were also identical to some hospital-based sequences, ruling out the possibility of completely excluding transmission from outside the hospital. However, the most parsimonious explanation is nosocomial transmission in those cases. Indeed, the subjects’ samples were collected at a time in Brazil when tests for SARS-CoV-2 infection were not easily accessible, and inpatients and HCW had to wait several days for a test result, thus presenting a risk for further transmission.

Single nucleotide variations were found across the entire SARS-CoV-2 genome. The spike (S) D614G mutation, found in all samples analyzed, has been associated with higher viral titers, suggesting increased viral infectivity [17]. Other variations were also found in different regions of the spike protein, including a 12-bp in-frame deletion that harbors part of the signal peptide and the predicted cleavage site in the beginning of S. As expected, the P323L change in the RNA-dependent RNA polymerase (RdRp), genetically linked to D614G, was also found in all our samples. *In silico* analysis showed that P323L may have an impact on the protein secondary structure, leading to a reduction in its molecular flexibility [19]. However, the phenotypic impact of these mutations is still poorly understood. Numerous other missense mutations were found that warrant further investigation concerning their phenotypes.

The most striking observation of our intrahost quasispecies variation analysis was that cancer patients carried significantly higher numbers of minor variants when compared to non-cancer counterparts (Fig 5). This difference was independent of, and unrelated to the Ct values obtained at the diagnostic tests, which did not differ between the two groups. Ct values (as a proxy to viral load) not only did not inversely correlate with virus diversity, but indeed showed a direct correlation, albeit with a low r_s_ coefficient. It is well established that naso- and oropharyngeal swabs are not the best types of sample for detecting SARS-CoV-2, compared to sputum for example, which contain a larger amount of viral genetic material [20]. Our data underscore the possibility that the variation in the viral quasispecies that we see is not generated in the naso- or oropharynx, but rather more distally in the respiratory tract (lungs) or even in other tissues such as the gut. Reports on the comparative expression of the virus’ cellular receptor ACE2 support the idea that those other tissues might be relevant sources of viral replication and, consequently, sites where diversity emerges [21, 22].

Unexpectedly, the intrahost quasispecies variation observed in cancer patients was not related to disease severity (requirement for ICU, death by any cause or COVID-19-related) or to the use of corticosteroids (which could lower their immunity status). Diversity was neither related to the use of oseltamivir, which was used by some patients to overcome a potential H1N1 infection until the COVID-19 diagnosis was released. Finally, the genetic diversity was not associated with the type of primary malignancy developed by the patient (solid tumor vs. hematologic tumors). Despite conflicting data existing in the literature, the hematologic cancer patients infected with SARS-CoV-2 herein analyzed did not show an increased chance of COVID-19 severe outcomes when compared to those with solid tumors [23].

Despite the fact we do not know the mechanisms by which or the anatomical sites where the SARS-CoV-2 quasispecies variation is generated in cancer patients, such increased variation compared to non-cancer patients may explain, at least in part, the more adverse outcomes to which cancer patients with COVID-19 are subjected to. By generating a higher number of distinct variants, the virus can explore wider areas of the sequence landscape and test variants with different regulatory and structural changes. Variation may impact tissue tropism, protein expression and function, stability, immune escape, drug resistance and pathogenicity. Further studies on SARS-CoV-2 diversity, especially in vulnerable patients with underlying comorbidities will shed light on our understanding of the underlying wide spectrum of disease outcomes associated with COVID-19 in humans.

## Methods

### Study population

Fifty-seven cancer patients followed at the Brazilian National Cancer Institute (INCA), Rio de Janeiro, Brazil, and 14 healthcare workers (HCW) diagnosed with COVID-19 between April 7^th^ and May 5^th^ 2020, early in the COVID-19 pandemic in Rio de Janeiro, were included in this study. SARS-CoV-2 infection was diagnosed through naso- and oropharyngeal swab specimens using RT-qPCR following the U.S. Centers for Disease Control and Prevention (CDC) protocol [24].

### Ethics Statement

All participants agreed to be enrolled in the study and signed an informed consent. Participants’ data were treated anonymously. This study was approved by the Brazilian National Commission for Ethics in Research (CONEP) (approval number: CAAE 30608220.8.0000.5274).

### SARS-CoV-2 nucleic acid isolation, amplification and sequencing

Naso- and oropharyngeal swabs were collected and placed into a conical tube containing 2 ml of viral transport medium (VTM, Thermo Fisher Scientific, Waltham, MA). Viral DNA and RNA were extracted with the QIAamp MiniElute Virus Spin Kit (QIAGEN, Chatsworth, CA) according to manufacturer’s instructions. All cDNAs were synthesized in duplicate using the SuperScript™ III First-Strand Synthesis System (Thermo Fisher Scientific). The SARS-CoV-2 complete genome amplification was based on an openly available protocol developed by the ARTIC network (https://artic.network/ncov-2019, accessed March 26, 2020) using the V.3 multiplex primers scheme and Platinum Taq DNA Polymerase High Fidelity (Thermo Fisher Scientific). Positive PCR products were purified with the ReliaPrep™ DNA Clean-Up and Concentration System (Promega, Madison, WI). Genomic libraries were constructed with the Nextera XT DNA Sample Preparation kit (Illumina Inc., San Diego, CA) according to the manufacturer’s protocol, pooled with 1% denatured PhiX DNA (sequencing control) and sequenced in a MiSeq platform (2× 251 cycles paired-end run; Illumina). New PCR reactions using combinations of the primers described above were carried out to cover regions with low coverage for each sample. Positive products were purified and sequenced by Sanger using the *BigDye Terminator kit* (Thermo Fisher Scientific) in an automated 3130XL Genetic Analyzer (Thermo Fisher Scientific). Sequences were edited and assembled with SeqMan v.7.0.0 (DNAStar Inc., Madison, WI).

### SARS-CoV-2 near full-length consensus sequence and nucleotide variations

All analyses were conducted using Geneious R11 software (Biomatters, Auckland, New Zealand), where the reads were trimmed to achieve an error rate below 0.1% and assembled to the Wuhan-Hu-1 reference sequence genome (GenBank number MN908947). A minimum mapping quality of 30 was required, providing a 99.9% confidence level that the mapping is correct. Additionally, all assemblies were visually inspected to evaluate the mapped reads and consequently to ensure the quality of the consensus generated and single nucleotide variation (SNVs) analysis. Consensus sequences representing SARS-CoV-2 near full-length genomes were extracted for each sample and aligned to the Wuhan-Hu-1 reference sequence genome. Nucleotide variations in relation to the reference sequence were identified and classified as SNVs. Intrahost single nucleotide variation (iSNV) was defined as a variation with a frequency greater than 2% and depth coverage by at least 500 reads. iSNVs were manually verified, and the intrahost viral genetic diversity rate was calculated as the number of nucleotide substitutions with a frequency greater than 2% for the given sample divided by the number of positions with depth coverage greater than 500 times multiplied by 10^−4^ (substitutions/site × 10^−4^).

### SARS-CoV-2 classification and phylogenetic analysis

For SARS-CoV-2 lineage classification, consensus genomes were submitted to *Pangolin* software (https://github.com/cov-lineages/pangolin, downloaded on June 10^th^, 2020) and to *CoV-GLUE* lineage system (http://cov-glue.cvr.gla.ac.uk/#/home, accessed on June 10^th^, 2020)[25], both based on the nomenclature proposed by Rambaut et al [26]. An alignment including the consensus sequences generated and genomes from Brazilian sequences available on the GISAID Database classified as B1, B1.1 and the Brazilian clusters B1.1-BR/ B1.1-EU/BR (S1 Table) were submitted to a maximum likelihood phylogenic reconstruction using PhyML v.3.0 and the best model of nucleotide substitution was defined with Model Generator (GTR) to investigate the sublineage classification of the study sequences [16, 27, 28]. Furthermore, a phylogenetic analysis that included the generated consensus sequences generated along with all SARS-CoV-2 sequences from Rio de Janeiro state (Brazil) presently available at GISAID (https://www.epicov.org/epi3/frontend, accessed on July 27^th^, 2020, S1 Table) was performed in order to investigate epidemiological relatedness of sequences.

### Statistical analyses

Mann-Whitney two-tailed test was used to compare intrahost diversity (substitutions/site × 10^−4^) between cancer patients and HCWs and between cancer patients’ clinical categorical variables. Spearman’s rank was employed to evaluate the correlation between intrahost diversity and continuous variables (such as age and SARS-CoV-2 RT-qPCR Ct values). All graphical representations and statistical analyses were performed using Geneious R11 (Biomatters) and GraphPad Prism v.8.0.1 (GraphPad Software Inc., San Diego, CA).

## Acknowledgements

We would like to thank all the participants of the INCA COVID-19 Task Force, clinical staff and patients from the Brazilian National Cancer Institute (INCA) for providing conditions and samples that enabled the conduction of this study. We also thank PhD. Renata Olício for providing support to Sanger DNA sequencing. We kindly acknowledge GISAID Database (https://www.gisaid.org/), the authors and laboratories for the SARS-CoV-2 genomes data shared. A table containing all genome sequences used in this article and the respective information can be found on S1 Table.

INCA COVID-19 Task Force: Luiza M. Abdo, Maria Theresa Accioly, Lucas R. Almendra, Rodrigo O.C. Araujo, Elisa Bouret C. Barroso, Marcelo A. Bello, Anke Bergmann, Ricardo S. Bigni, Martin H. Bonamino, Franz S. Campos, Samuel Z.B. Cordeiro, Susanne Crocamo, Magda S. da Conceição, Jesse L. da Silva, Carolina S. Dantas, Lucas Z. de Albuquerque, Roberto R.M. de Araujo Lima, Renata de Freitas, Fernando L. Dias, Jorge L.A. Dias, Michelle M.Q. dos Santos, André F. Duarte, Sima E. Ferman, Vanessa C. Fernandes, Erico L. Ferreira, Priscila S. Ferreira, Kelly M. Fireman, Carolina Furtado, Marianne M. Garrido, Renan G. Gomes Junior, Bruno A.A. Gonçalves, Juliana G. Gonçalves, Gustavo H.C. Guimarães, Nelson J. Jabour Fiod, Ana Cristina M. Leão, Décio Lerner, Valdirene S. Lima, Eduardo Linhares, Monique S.A. Lopes, Ianick S. Martins, Bruna P. Matta, Amanda S. Medeiros, Ana Cristina P. Mendes Pereira, Paulo A. Mora, Miguel A.M. Moreira, Daniela P. Oliveira, Alexandre M. Palladino, Diego J.G. Paula, Ana C. Pecego, Bárbara C. Peixoto, Patrícia A. Possik, Gelcio L. Quintella Mendes, Matheus A. Rajão, Maria D.M. Rocha, Fernando L.B. Rocha Gutierrez, Luciana de O.R. Rodrigues, Giovani B. Santos, Marcelo R. Schirmer, Karina L. Silva, Lilian S. Silva, Antonio A.D. Souto, Leandro S. Thiago, Luiz C.S. Thuler, Fabiana Tonellotto, Gisele M. Vasconcelos, Dolival L. Veras Filho.

## Author contributions

JDS, LRG, BMA and MAS conceived and designed the study. JDS, LRG and BMA optimized all reagents and performed the sequencing and analysis. ACM collected clinical data. CC, JA, JPBV, ACM and MAS provide expert advice on experimental planning and data interpretation. JDS, LRG, BMA and MAS wrote the manuscript. JDS, LRG, BMA, CC, JA, JPBV, ACM and MAS revised and edited the manuscript.

## Supporting information

**S1 Fig. Depth coverage across SARS-CoV-2 genome.** Samples’ depth coverage is shown in gray and median coverage in red. Genome coordinates are relative to the SARS-CoV-2 Wuhan-Hu-1 reference sequence genome (GenBank acc.# MN908947).

**S2 Fig. Spearman correlation analysis of viral genetic diversity and patients’ age.** No correlation was observed between viral genetic diversity and patients’ age. Spearman correlation analysis r_s_ and p-values are indicated.

**S3 Fig. Viral genetic diversity association with clinical outcomes.** Tukey boxplots show viral genetic diversity distribution according to the following clinical criteria: death (A), death from COVID-19 (B), admission to intensive care unit (ICU) (C), use of corticosteroid (chronic or during COVID-19 diagnostics) (D), use of oseltamivir (during COVID-19 diagnostics) (E), type of cancer (hematological vs. solid tumors) (F). All comparisons were submitted to Mann-Whitney test (two-tailed), no statistically significant differences were found.

**S1 Table. SARS-CoV-2 sequences downloaded from GISAID database.**

**S2 Table. Synonymous and non-synonymous nucleotide variations found in the 71 samples analyzed.**

